# Establishment of an intragastric surgical model using C57BL/6 mice to study the vaccine efficacy of OMV-based immunogens against *Helicobacter pylori*

**DOI:** 10.1101/2023.11.16.567319

**Authors:** Sanjib Das, Prolay Halder, Soumalya Banerjee, Asish Kumar Mukhopadhyay, Shanta Dutta, Hemanta Koley

## Abstract

Chronic gastritis is one of the major symptoms of gastro-duodenal disorders typically induced by *Helicobacter pylori* (*H. pylori).* To date, no suitable model is available to study pathophysiology and therapeutic measures accurately. Here, we have presented a successful surgical infection model of *H. pylori-*induced gastritis in C57BL/6 mice that resembles features similar to human infection. The proposed model does not require any preparatory treatment other than surgical intervention. C57BL/6 mice were injected with wild-type SS1 (Sydney strain 1, reference strain) directly into the stomach. Seven days post infection, infected animals showed alterations in cytokine responses along with inflammatory cell infiltration in the lamina propria, depicting a prominent inflammatory response due to infection. To understand the immunogenicity and protective efficacy, the mice were immunized with outer membrane vesicles (OMVs) isolated from an indigenous strain with putative virulence factors of *H. pylori* [A61C (1), *cag+/vacA s1m1*]. In contrast to the nonimmunized cohort, the OMV-immunized cohort showed a gradual increase in serum immunoglobulin(s) levels on the 35^th^ day after the first immunization. This conferred protective immunity against subsequent challenge with the reference strain (SS1). Direct inoculation of *H. pylori* into the stomach influenced infection in a short time and, more importantly, in a dose-dependent manner, indicating the usefulness of the developed model for pathophysiology, therapeutic and prophylactic studies.

## 1. Background

Gastroduodenal disorders are the cumulative effect of carefully orchestrated molecular interactions between host and pathogen factors belonging to the genus *Helicobacter* [1]. With almost 50% of the population worldwide infected by the pathogen, it is one of the major health burdens in developing nations [2]. Although *H. pylori* has been recognized as a class I carcinogen by the WHO, very little has been explored thus far. This is primarily due to the asymptomatic nature of infected individuals, expensive clinical detection (e.g., endoscopy, urea breath test, etc.) and diagnosis with considerable information scarcity [3]. In addition, the global antimicrobial resistance (AMR) pattern of *H. pylori* is changing alarmingly, resulting in a paradigm shift in “treatment of choice” by clinical practitioners [4].

Research in *in vitro* and *in vivo* systems of *H. pylori* is continuously enriching our understanding of pathophysiology and genetic predisposition related to adaptation, survival and coevolution of the pathogen [5,6]. For instance, *H. pylori* has the inherent ability to modulate the gastric microenvironment, such as increasing gastric pH by means of urease upregulation, employing different adhesion proteins or simply dislodging itself when the pH becomes overwhelmingly acidic [7,8]. Such responses, along with others, act as the precursor to a chronic infection that largely depends on the gastric acid neutralizing capacity unique to each strain. The pathogen is known to recruit different adhesins depending upon the stages of disease progression, such as BabA during early infection or SabA during ongoing inflammation [9]. In addition, host antigens present on the surface of host cells, mucins and other gastric cells, such as A/B-Le^b,^ MUC5AC, MUC1 and H type 1, play important roles in bacterial adhesion, further promoting the severity of different gastric maladies [10,11].

To date, a combination of antibiotics with a proton pump inhibitor (PPI) is the only mode of treatment available due to the lack of a potent vaccine [12]. Moreover, an efficient animal model is crucial to understanding the immunological attributes of different immunogen(s) for vaccine development, which existing models fail to satisfy. Considerable efforts have been made to establish a reliable murine (gerbil or mouse) model to serve this purpose, including extensive application of transgenic animals with single or double mutations, but unfortunately, no significant efforts have been made toward the route of administration to induce an infection [13]. The preexisting method relies on the oral administration of multiple doses of inoculums along with antibiotic pretreatment to induce an infection [14]. In addition, it takes a minimum of two weeks to develop an infection using the traditional approach, which is significantly higher than any other enteric pathogens, such as *E. coli* or *Salmonella,* while using an animal model [15–17].

Therefore, in this study, we introduced an infection by surgically exposing the stomach of C57BL/6 mice by directly injecting *H. pylori* inoculums. We assessed different immunological markers for active infection and applied the same to study the vaccine efficacy of OMV-based immunogens isolated from a prevalent strain.

## 2. Methods

### 2.1. Bacterial strains and culture conditions

Both SS1 and A61C (1) (clinical strain) were revived from glycerol stock using brain heart infusion agar (BD Difco, USA) supplemented with 7% horse blood, 0.4% IsoVitaleX with antibiotics such as amphotericin B, trimethoprim, and vancomycin (Sigma Aldrich, USA) at concentrations as described previously [18]. Inoculated plates were then kept under microaerophilic conditions (5% O2, 10% CO2, and 85% N2 at 37°C) for 48 hrs and sub-cultured before conducting any experiment.

Broth culture was prepared using Brucella Broth (BB broth) (BD, Difco, USA) supplemented with 10% HI-horse serum and vancomycin (Sigma, USA). The inoculated flask was then kept in shaking conditions (100 rpm) overnight while maintaining the microaerophilic environment [19].

### 2.2. Characterization and selection of strains

All strains were checked for oxidase, catalase and urease as mentioned elsewhere [20]. Next, an antibiogram was performed using the agar dilution method following CLSI guidelines (**supplementary 1**). PCR-based detection was applied for genotypic characterization. Some major virulence factors, such as *cagA, vacA, babA* and *dupA,* were checked using either simplex or multiplex PCR [21,22]. The primers used in the present study are listed in **supplementary 2**.

### 2.3. Animals

Six- to eight-week-old female C57BL/6 mice were received from the NICED-Animal house facility. The animals were kept in specific pathogen-free (SPF) conditions maintained at 25±°C with 65 ± 2% humidity and a 12/12-hour light/dark cycle. Animals weighing ∼23 grams were selected for the study and provided with sterile food and water ad libitum. All experiments were performed following the standard operating procedure outlined by the Committee for the Purpose of Control and Supervision of Experiments on Animals (CPCSEA), Ministry of Environment and Forest, Government of India. The Institutional Animal Ethics Committee (IAEC) of NICED with registration no. 68/GO/ReBi/S/1999/CPCSEA valid 17.07.2024 approved (Approval No. PRO/194/June 2022-25) and supervised experimental design and protocols from time to time.

### 2.4. Animal experimental design

Thirty-six female C57BL/6 mice were randomly assigned into two major groups, each comprising 18 animals. To determine an infectious dose for the surgical model, the 1^st^ set of mice was further divided into three subgroups and infected with a dose of either 1x10^8^ (n=6) or 2x10^8^ CFU/mL (n=6) along with PBS control (n=6). All groups were housed for one or two weeks under sterile conditions.

For immunological studies, the remaining 18 mice were separated into two groups: nonimmunized (NI) (n=6) and oral or i.p. immunized (IM) (n=6 in each group). An oral or intraperitoneal immunization with 50 μg of OMVs dissolved in PBS was administered on days 0, 14, and 28. Blood was collected at different time points, and the serum was isolated and stored at -20°C for use in different immunological assays. For the protective efficacy study, both groups (IM, NI) were infected surgically on the 35^th^ day post-first immunization and sacrificed 7 days post-infection **(Figure 1)**.

**Fig 1:**
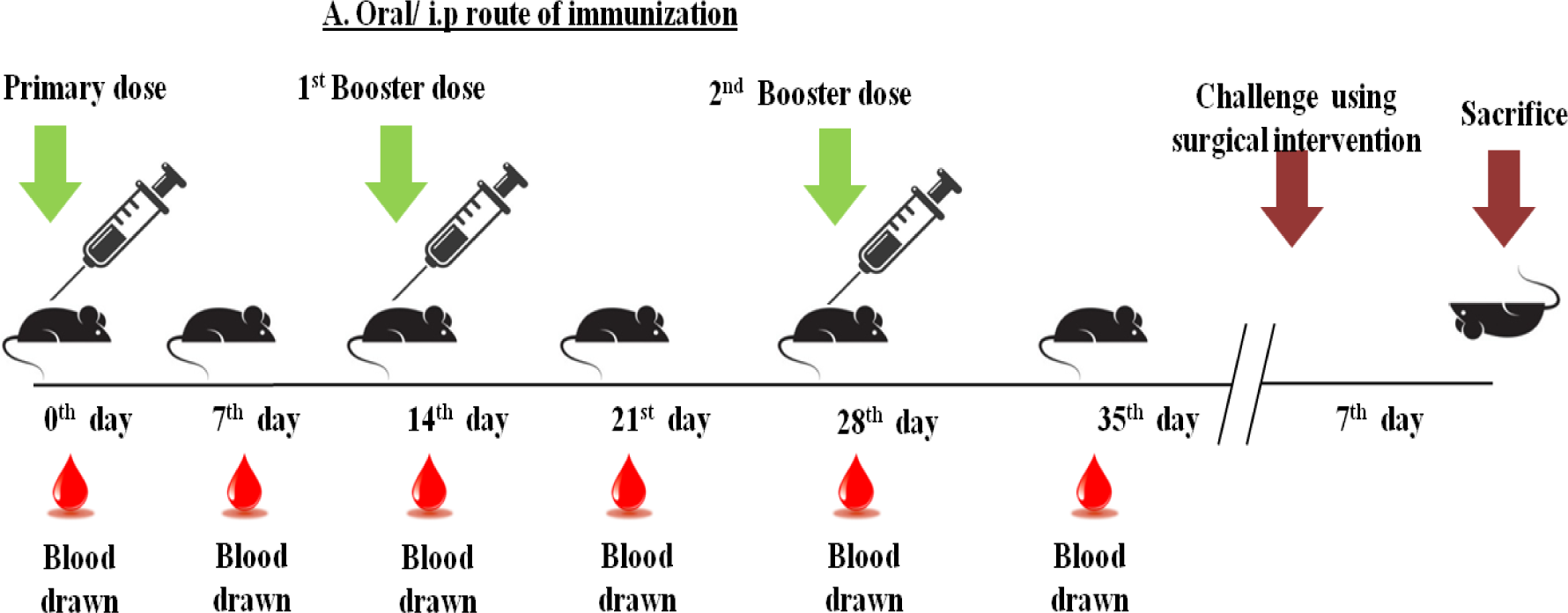
Immunization and blood collection schedule

### 2.5. Surgical model development

Experimental animals were kept in fasting conditions overnight with sterile water. Initially, animals were sedated by an intraperitoneal injection of a mixture of ketamine (87.5 mg/kg) and xylazine (12.5 mg/kg) [23]. The stomach was exposed through a 2-3 cm midline incision without compromising any major blood supply. A disposable syringe with a 26G needle containing 200 μl (∼2x10^8^ CFU/mL) of the inoculums in PBS was directly injected into the stomach. Hydration was maintained in the exposed stomach using sterile normal saline (0.9%) throughout the surgery. The stomach was placed back inside the abdominal cavity, and the incision was sutured back. The incision site was monitored for any infection and occasionally washed with 5% povidone-iodine (betadine) soaked in a sterile gauge for 72 hours [24]. Sterile food and water were provided to the animals once they regained consciousness (**Figure 2**).

**Fig 2:**
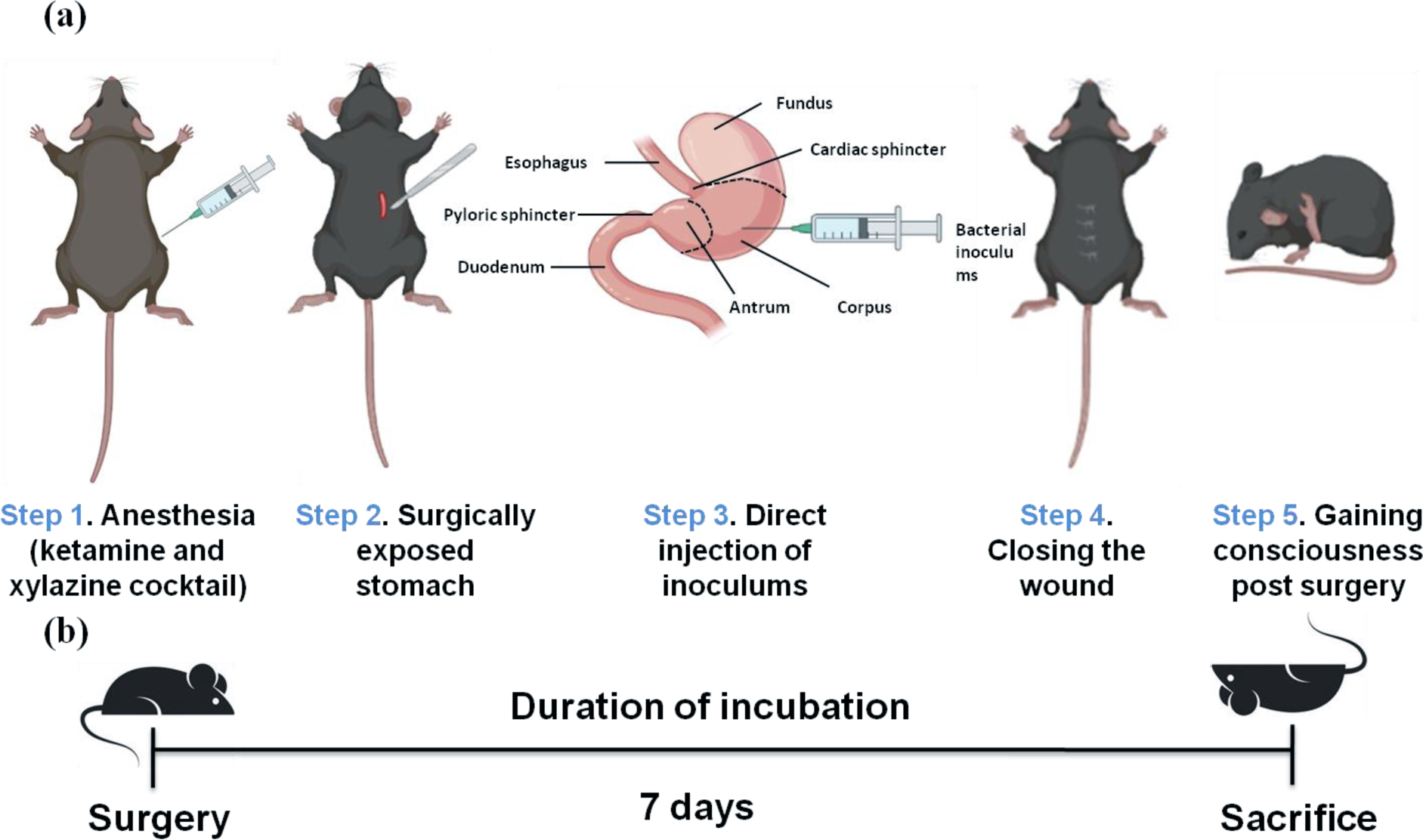
(a) Graphical Representation of the “Surgical Model” using C57BL/6.Bacterial 8 inoculation (∼2x10 CFU/mL) is directly injected into multiple sites of the stomach, **(b)** Schematic diagram from infection to sacrifice

### 2.6. Post-OP observation

All infected mice were observed twice a day for 7 days. Physical parameters were checked along with stool consistency and the nature of mucus or blood (if any) present in the feces. Rectal swabs were taken daily and were subjected to RUT solution and spread-plate to observe the shedding of the organism. Isolated colonies (if any) were confirmed using a PCR-based technique.

1. *H. pylori* infection augments the modulation of both pro- and anti-inflammatory cytokines in the host [25]. Therefore, IL-1β, TNFα, IFNγ, IL-6, IL-10 and IL-17 were tested using cytokine measuring kits (Invitrogen, USA) following the manufacturer’s protocol. Fifty microliters of serum samples from uninfected, 7-day- and 14-day-infected mice were used to quantify the inflammatory response due to infection.

### 2.7. Immunogen preparation

Outer membrane vesicles (OMVs) were isolated from the *Helicobacter* strain [A61C (1), *cagA+,vacAs1m1*] following the methods described by Bhaumik et al., 2023, with slight modification [26]. In brief, BB broth (BD, Difco, USA) was inoculated with log phase (OD600 ∼0.6) preculture of the respective strains and kept overnight in microaerophilic conditions under constant shaking (120 rpm) at 37°C. On the next day, centrifugation was performed consecutively first at 8000xg for 15 min, followed by 30 min. The supernatants were then filtered twice with 0.45 μm and 0.22 μm syringe filters (Millipore, USA). To prevent protein degradation, a protease inhibitor cocktail was incorporated into the filtrate and ultra-centrifuged at 140,000 x g at 4°C for 4 hrs using a P27A-1004 rotor (Hitachi). A density gradient centrifugation allowed obtaining the purified OMVs. Protein content was measured using a Lowry protein estimation kit (Pierce, USA) and stored at -80°C until further use.

### 2.8. Characterization of the Immunogen

#### 2.8.1. Dynamic Light Scattering and Zeta potential

Concentrated OMVs were diluted 10-fold to reach a concentration of 0.1 mg/mL. The hydrodynamic size of OMVs was measured using a Malvern Zetasizer ZS90 (Malvern Instruments, Germany) and analyzed using ZS Xplorer version 3.1.0.64 [27].

#### 2.8.2. Transmission electron microscopy

Diluted OMVs were placed on a carbon-coated grid and left for 10-20 min for absorption. The samples were then washed twice with drops of Tris buffer solution. Excess fluid was soaked using blotting paper, followed by staining with 2% uranyl acetate and air drying. Finally, the OMV- coated grids were observed under a JEOL JEM 2100 HR (JEOL, Tokyo, Japan) [28].

#### 2.8.3. LC/MS of OMVs and analysis

Proteins present in OMVs were used for digestion and reduced with 5 mM TCEP and further alkylated with 50 mM iodoacetamide and then digested with trypsin (1:50, trypsin/lysate ratio) for 16 h at 37°C. Digests were cleaned using a C18 silica cartridge to remove the salt and dried using a speed vac. The dried pellet was resuspended in buffer A (2% acetonitrile, 0.1% formic acid).

Experiments were performed on an Easy-nlc-1000 system coupled with an Orbitrap Exploris mass spectrometer. One microgram of peptide sample was loaded on a C18 column (15 cm, 3.0 μm Acclaim PepMap, Thermo Fisher Scientific), separated with a 0–40% gradient of buffer B (80% acetonitrile, 0.1% formic acid at a flow rate of 500 nl/min) and injected for MS analysis. LC gradients were run for 60 minutes. MS1 spectra were acquired in the Orbitrap (Max IT = 25 ms, AGQ target = 300%; RF lens = 70%; R=60 K, mass range = 375−1500; profile data). Dynamic exclusion was employed for 30 s, excluding all charge states for a given precursor. MS2 spectra were collected for the top 12 peptides. MS2 (Max IT= 22 ms, R= 15 K, AGC target 200%). All samples were processed, and the generated RAW files were analyzed with Proteome Discoverer (v2.5) against the UniProt organism database. For dual Sequest and Amanda searches, the precursor and fragment mass tolerances were set at 10 ppm and 0.02 Da, respectively. The protease used to generate peptides, i.e., Enzyme specificity was set for trypsin/P (cleavage at the C-terminus of “K/R: unless followed by “P”). Carbamidomethyl on cysteine as a fixed modification and oxidation of methionine and N-terminal acetylation were considered variable modifications for the database search.

### 2.9. Extraction of lipopolysaccharide (LPS) and outer membrane proteins (OMPs)

LPS and OMPs were extracted following the methods described by Mukherjee et al. (2016) [29]. LPS was then treated with proteinase K to ensure the absence of any protein residue. The carbohydrate content of LPS was then quantified using the phenol‒sulfuric acid method and measured at a wavelength of 492 nm [30]. Isolated proteins were quantified for their concentration using a Modified Lowry’s Kit (Pierce, USA) and measured at 660 nm using a spectrophotometer.

### 2.10. ELISA

Serum immunoglobulin (IgG, IgM, IgA) levels were measured against OMPs or LPS following the method of Howlader et al., 2018 [31]. Twofold serial dilutions were prepared from serum isolated from both immunized and nonimmunized groups. HRP-conjugated secondary anti-mouse IgG, anti-IgA, and anti-IgM antibodies (Sigma Aldrich, USA) were used to detect the antibody titer. Each experiment was replicated thrice with pooled sera from different groups.

### 2.11. Serum Bactericidal Assay and Scanning Electron Microscopy (SEM)

The effect of immunized mouse sera on bacterial morphology was measured and visualized using scanning electron microscopy (SEM) following a previously described protocol [32]. Bacteria along with heat-inactivated mouse sera and 25% guinea pig complement (with/without) were incubated for 1 hr under microaerophilic conditions followed by plating for viable colonies or fixation with 3% glutaraldehyde overnight followed by a gradual dehydration step initially with alcohol and then substitution later with a mixture of alcohol and hexamethyldisilazane (HMDS) at ratios of 2:1, 1:1 and 1:2. Finally, the samples were mounted on specimen stubs, sputter-coated with gold and analyzed on a Quanta 200 SEM (FEI, Netherlands).

### 2.12. Cytokine assay

Both immunized and nonimmunized mice were sacrificed, and the spleens were harvested. After isolating spleen cells, ∼10^5^ cells were incubated with 50 μg of OMVs and incubated overnight. IL- 10, IFN-γ, IL-1β, IL-6, IL-4, TNF-α and IL-17 were measured in the culture supernatant using a cytokine measuring kit (Invitrogen, USA) [33].

### 2.13. Fluorescence-activated cell sorting (FACS) analysis

Spleen cells were harvested and re-stimulated using isolated OMVs (50 μg) and incubated overnight. The next day, the cells were scraped, washed thoroughly, blocked and then incubated with mouse anti-CD4+, CD8+ or CD19+ antibodies. Unbound antibodies were washed, and a specific epitope of the immune cell population was observed using FACS Aria II [32].

### 2.14. Protective efficacy Study

Seven days after the last immunization, both the immunized and control groups were challenged with the wild-type SS1 strain using a newly developed surgical procedure and housed for 7 days before being sacrificed. The antrum was identified and separated into two parts. Half of each part was immediately transferred to BHI kept on ice, and the other half was transferred to neutral buffered formalin (NBF, 10%) solution to fix the tissue and left at room temperature. Harvested tissue in BHI was weighed and homogenized using a micro pestle and serially diluted using PBS. The diluted samples were then spread onto BHIA and kept under microaerophilic conditions for 3-5 days. Any visible colonies were then counted and confirmed using RUT and PCR. Histopathological assays were performed as described elsewhere [34,35]. Briefly, samples kept in 10% formalin were washed and gradually dehydrated using the alcohol gradation method followed by preparing a paraffin block. A thin section (approximately 5 μm) was prepared using a microtome. The slides were then dewaxed, rehydrated, and stained. Hematoxylin-eosin or Giemsa stains were used for the study because they enhance tissue or bacterial contrast. Finally, the slides were mounted and observed under a microscope. Histological scoring was assigned for each sample based on their morphological changes.

### 2.15. Statistical analysis

The presented data do not follow a normal distribution due to biological variations. Nonparametric tests were adopted for all data analyses. Triplicate data were expressed as the mean ± SD (standard deviation) using GraphPad Prism version 5.02. Two-way analysis of variance (ANOVA) or the Mann‒Whitney test (for animal data) was performed as per the requirements, and statistical significance was determined from the *P* values mentioned in the figure legends.

## 3. Results

### 3.1. Characterization of the *H. pylori strain* used in the study

A61C (1) is a type I clinical strain isolated from the endoscopic sample of a patient diagnosed with gastric cancer, and SS1 is a mouse-adapted strain. Both the clinical isolate and challenge strains were positive for the rapid urease test (RUT), catalase and oxidase. The status of genetic characterizations of all strains is listed in **Table 1**. A61C (1) was selected for immunogen preparation, while SS1 was used as a challenge strain in this study.

**Table 1:**
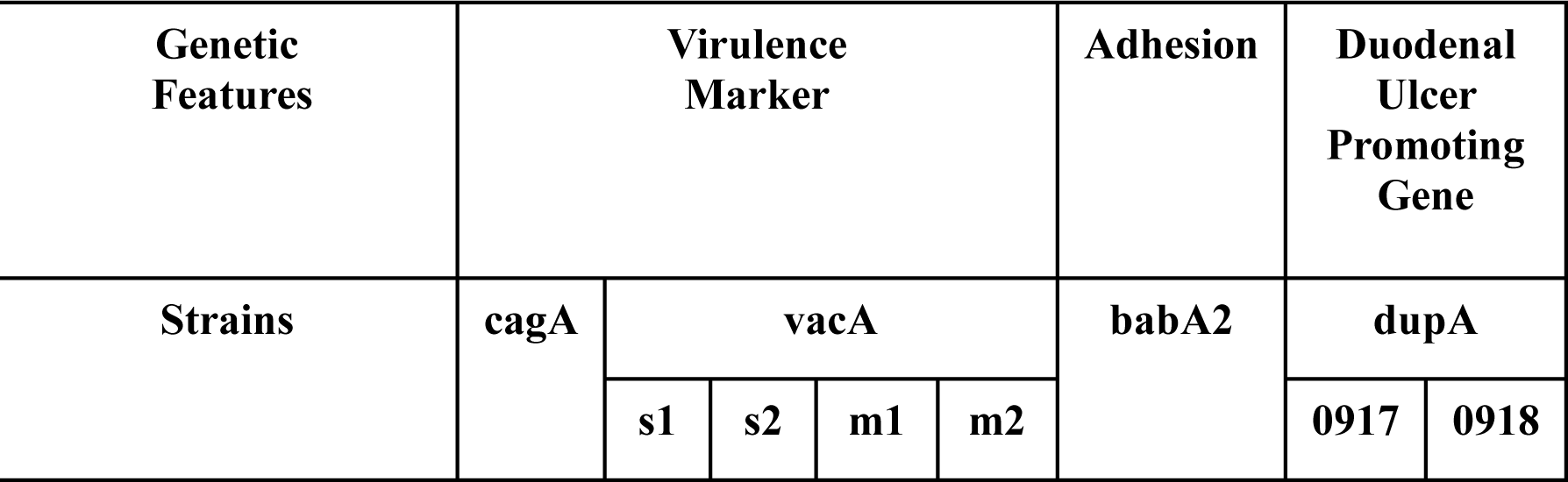
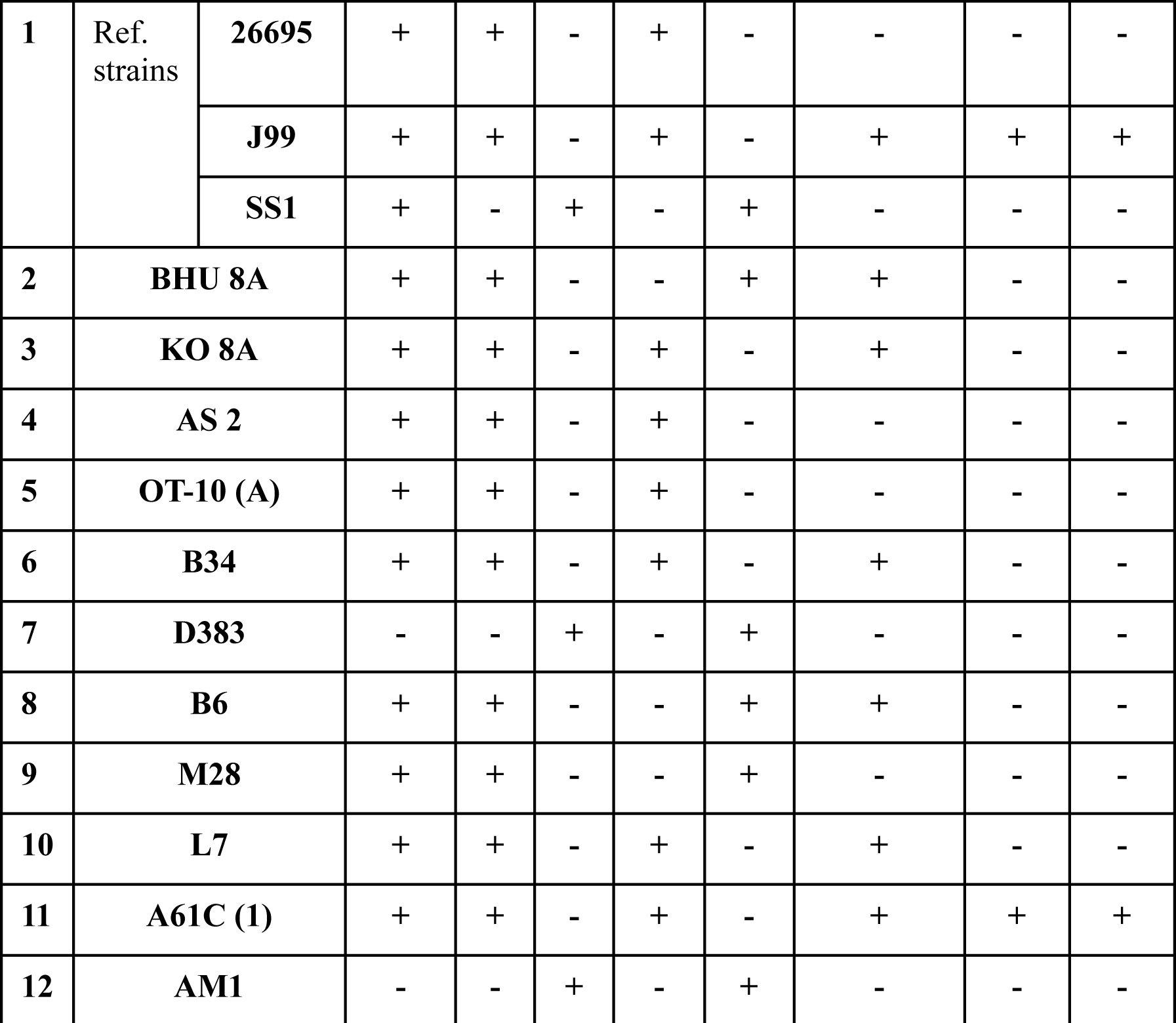
Major virulence factors.

### 3.2. Clinical response caused by surgical intervention

In the present study, 2x10^8^ CFU of bacteria were used to induce an active infection. Oral inoculation with the aforementioned dose revealed inconsistent results. Moreover, in the majority of cases, very little or no recovery of the bacterial population was observed using available detection techniques. Mice receiving WT SS1 directly to their stomachs by surgical means developed various degrees of gastric changes. Recovery of bacterial colonies from stool was insignificant and erroneous compared to gastric tissues, which were considerably higher (∼2-3 times) and were confirmed to be positive upon RUT, spread-plate and PCR. The recovery rate *of H. pylori* from the 7-day infected mice was comparatively higher than that at 14 days post infection (**Figure 3**).

**Fig. 3.**
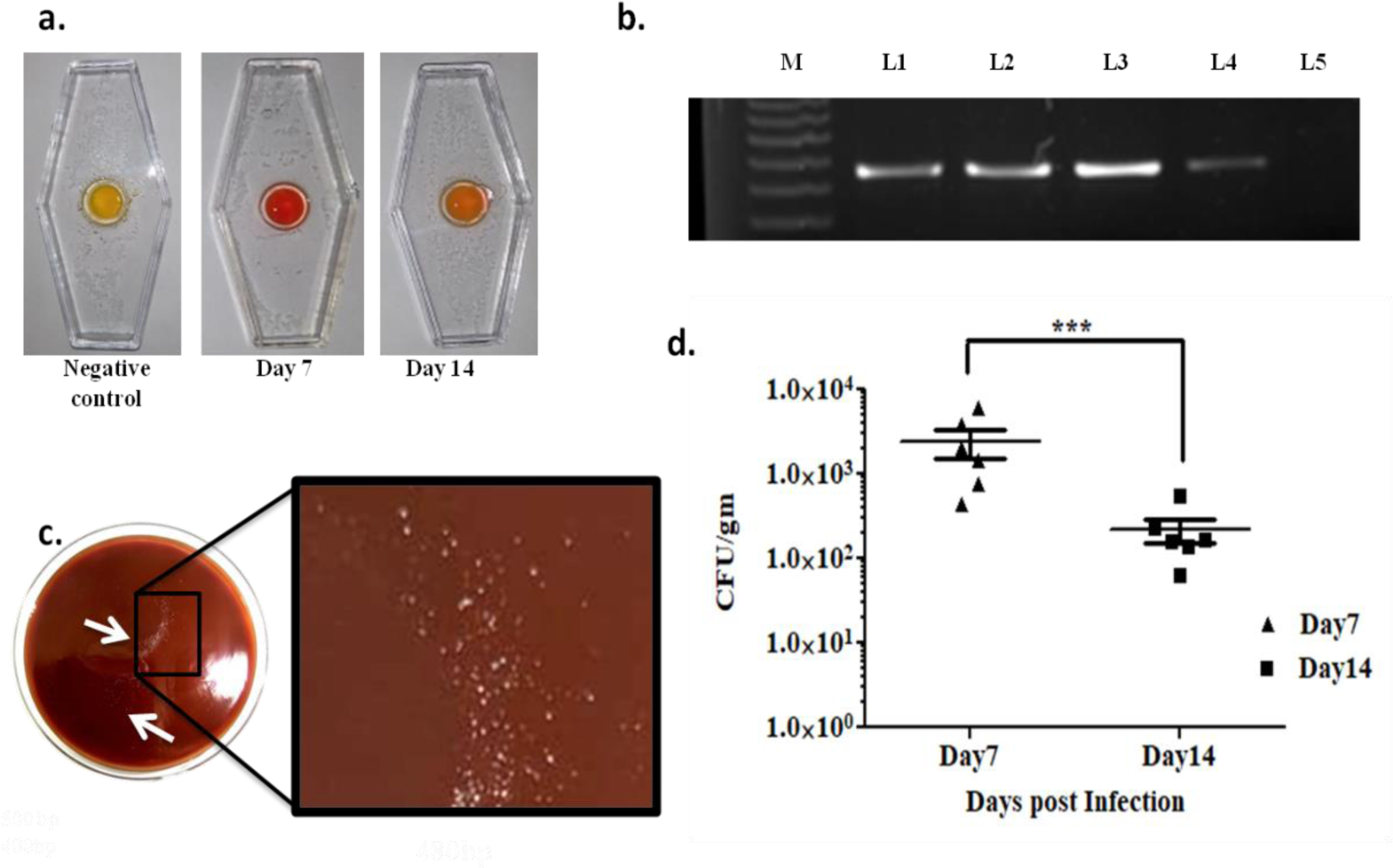
**a:** RUT of gastric tissue sample, b: Confirmatory ureB (480 bp) PCR for the presence of *H. pylori* recovered from the gastric tissue of infected mice. M-100 bp marker, L1 positive control, L2-L3 represent 7day post infection, L4 represents 14-day post infection, L5 negative control, c: *H. pylori* colonies recovered from gastric tissue **d.** Colonies recovered 7- and 14-days post infection, Data represented here are the mean values +/-Standard Deviations (SD) of three independent experiments. The differences in post infection day wise response of each of the studied cytokines were highly significant. Statistical significance was found between 0day 7day and 14day infected mice serum (***p<0.001)

### 3.3. Inflammatory response due to infection

Cytokine analysis of infected mice showed drastic differences in cytokine levels, such as IFNγ, IL-1β, TNF-α, IL-10, and IL-17. The majority of the pro-inflammatory cytokines were upregulated after 7 days post infection, except IL-6, which was found to be more pronounced at 14 days than at 7 days post infection (**Figure 4**).

**Fig. 4:**
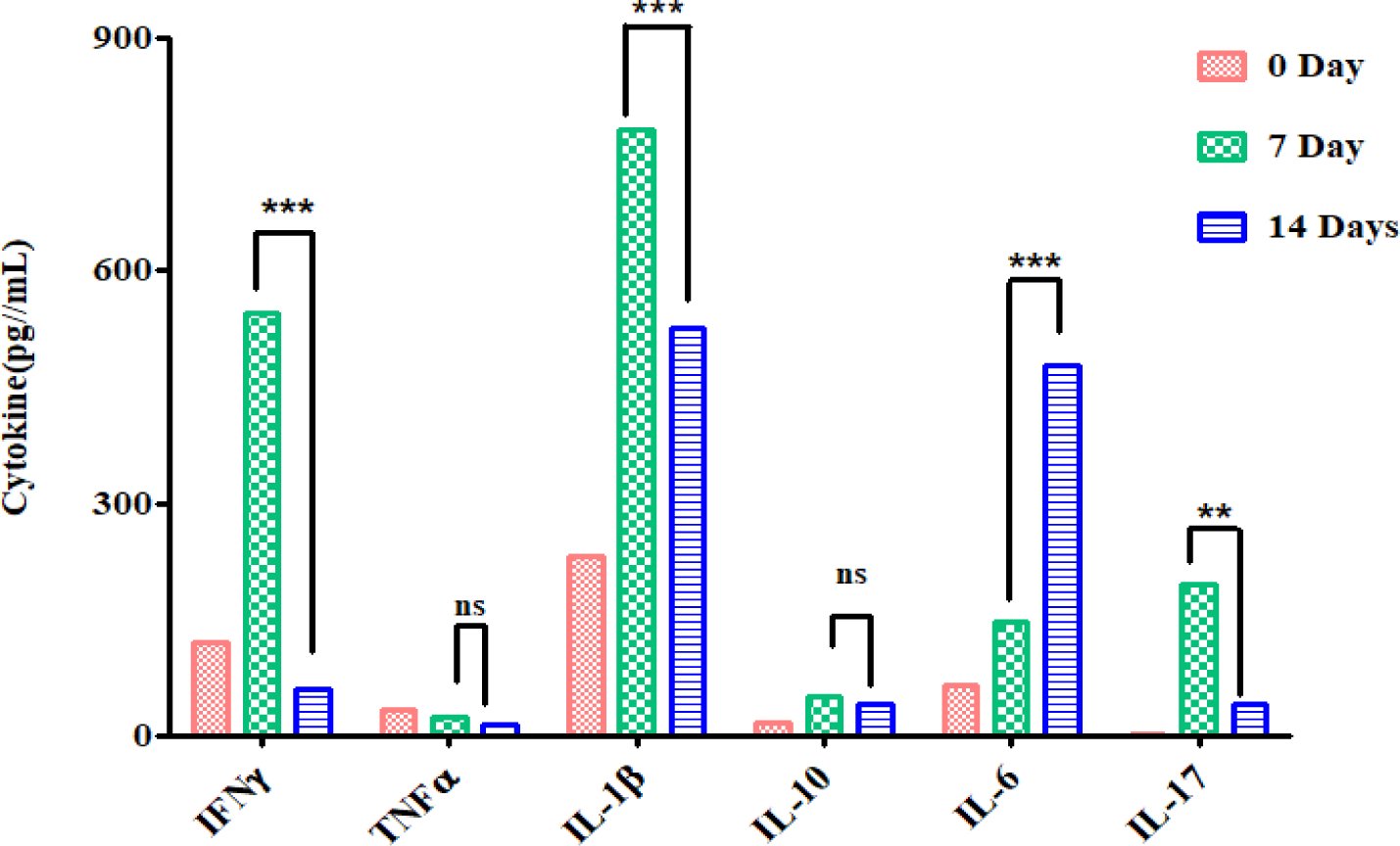
Histogram representing elevation of inflammatory cytokine levels on and 14-days post infection

### 3.4. Histopathological changes due to intragastric infection

The gastric tissue observed under a microscope revealed various degrees of gastritis, which was then categorized according to the Houston-updated Sydney System based on the infiltration of inflammatory cells within the lamina propria [36]. The changes were represented as Grade 0: absent inflammation, Grade 1: mild inflammation, Grade 2: moderate inflammation, and Grade 3: severe inflammation. The intensity of infiltration was greater on the 7^th^ day than on the 14^th^ day post infection. Structural changes were observed in the antrum of infected mice (**Figure 5**).

**Fig 5:**
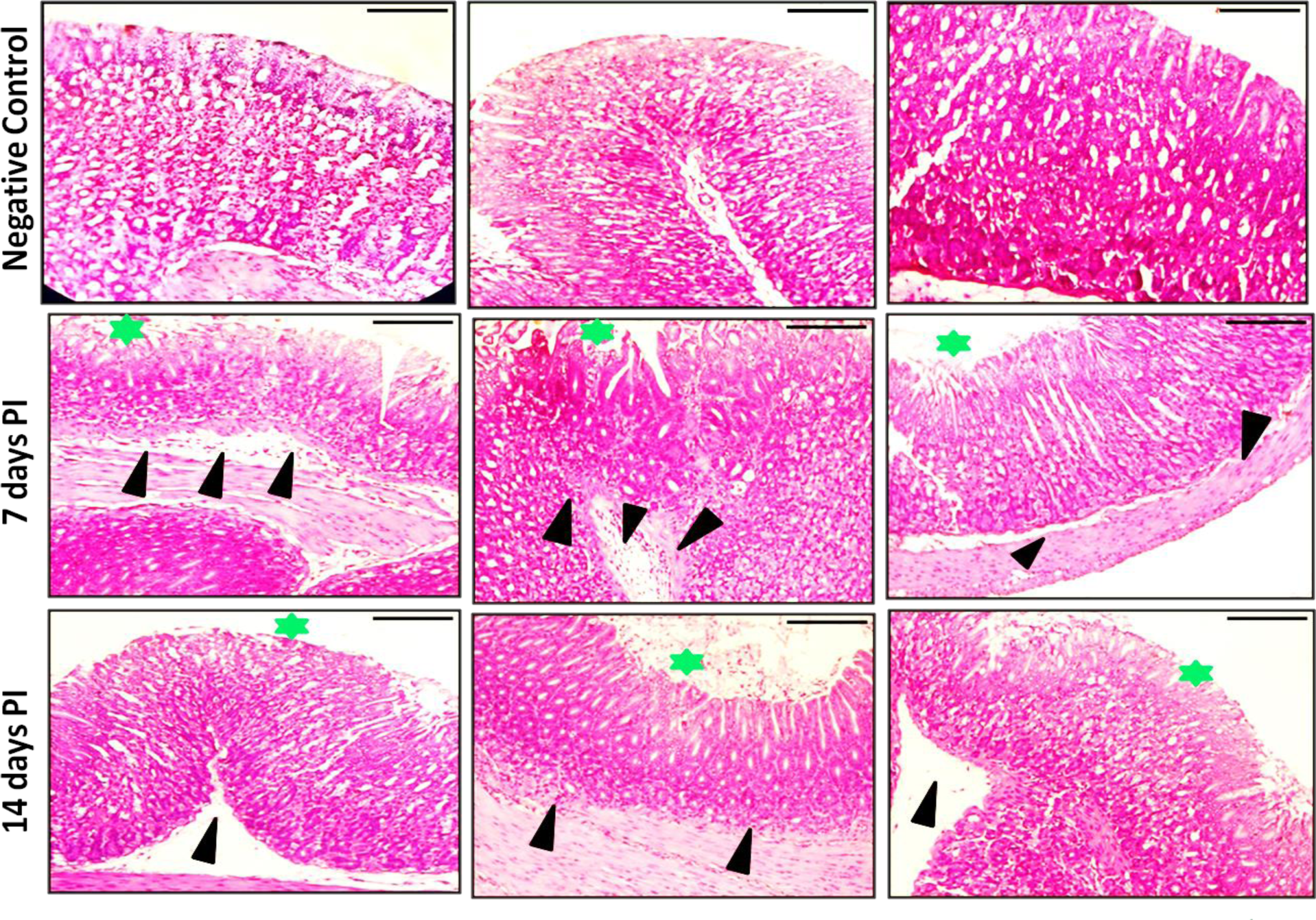
Histological observation of gastric epithelium, indicate mononuclear cell infiltration, 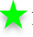 represent gastric epithelial lining. Changes observed in 7days and 14 days post infection (PI).

### 3.5. Isolation and characterization of *H. pylori* OMVs

Growth curve analysis (data not shown) revealed A61C (1) to be slow-growing compared to other clinical isolates with similar genetic backgrounds. Both SS1 and A61C (1) secrete OMVs during the log phase of growth. Dynamic light scattering revealed the size of OMVs with a mean peak of 50 nm. TEM images showed heterogeneity in OMVs, with the majority distribution ranging from 50-200 nm (**Figure 6**).

**Fig. 6:**
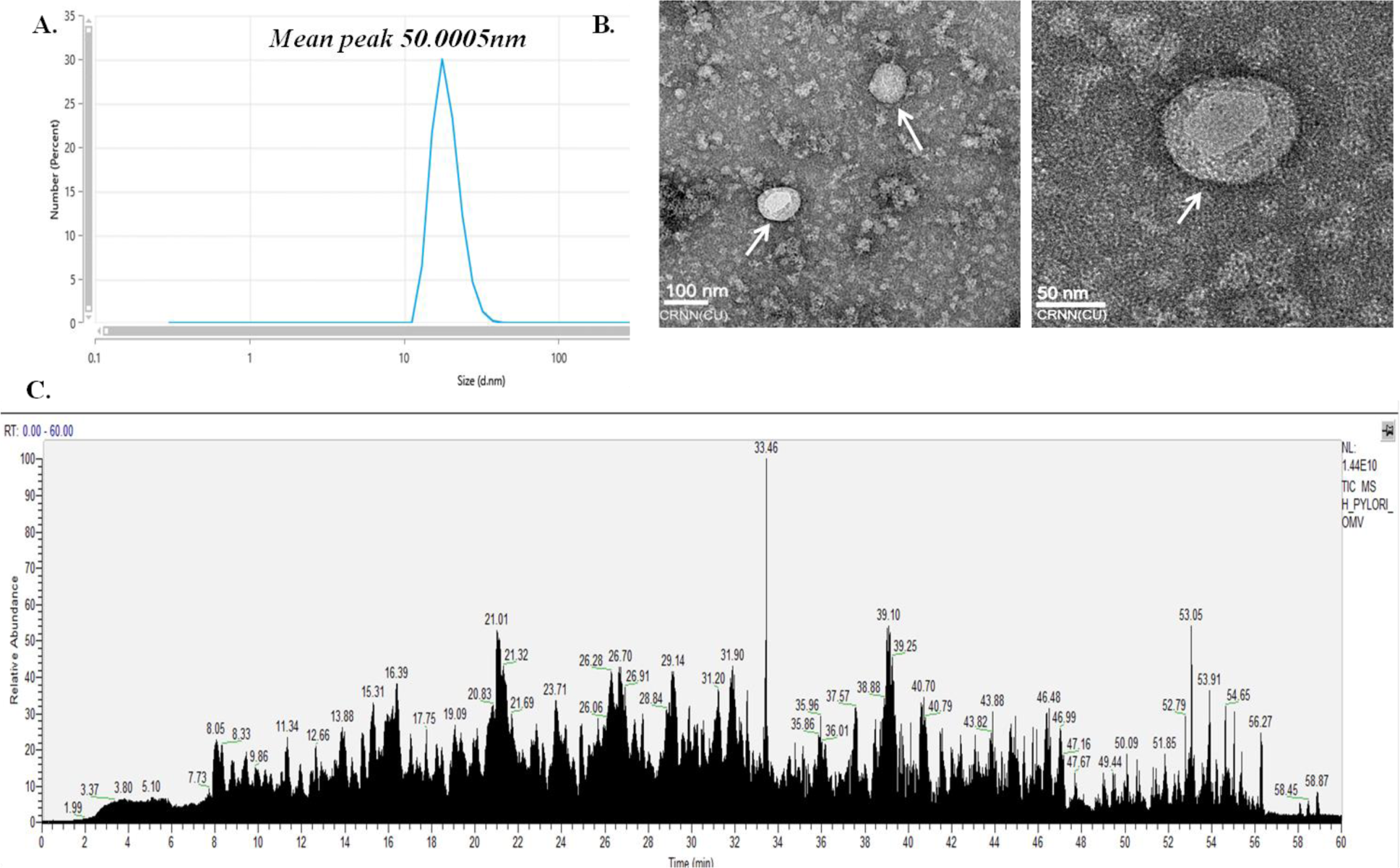
Biophysical characterization of OMVs; A. Dynamic Light Scattering showing a uniformity in OMVs population**, B.** Transmission Electron microscopy revealing the circular morphology of OMVs, **C.** Mass spectrometry analyses of OMVs

### 3.6. Proteomics of OMVs

The outer membrane vesicles of the selected strain to produce the immunogen consisted of different intracellular as well as outer membrane proteins that are listed in **Table 2**.

**Table 2:** Proteomics of OMVs.

| Accession | Protein | Molecular mass (kDa) | No. of peptides | Sequence coverage (%) |
| --- | --- | --- | --- | --- |
| P69996 | nickel cation binding, urease activity | 61.6 | 3 | 7 |
| P14916 | nickel cation binding, urease activity | 26.5 | 1 | 4 |
| P52093 | ferric iron binding, ferrous iron binding, ferroxidase activity | 19.3 | 2 | 19 |
| P42383 | ATP binding, ATP-dependent protein folding chaperone, isomerase activity, unfolded protein binding | 58.2 | 3 | 5 |
| O25723 | transmembrane transporter activity | 47.5 | 1 | 9 |
| O26011 | carboxy-lyase activity, flavin prenyltransferase activity | 20.6 | 1 | 18 |
| P56003 | GTP binding, GTPase activity, guanosine tetraphosphate binding, translation elongation factor activity | 43.6 | 2 | 4 |
| O25475 | ABC-type protein transporter activity, ATP binding, metal ion binding, protein-exporting ATPase activity | 99 | 1 | 3 |
| O24949 | Hypothetical protein | 23.6 | 1 | 3 |
| O26012 | Hypothetical protein | 24.3 | 1 | 3 |
| O25206 | ATP binding, DNA binding | 98.4 | 1 | 3 |
| P56035 | rRNA binding, structural constituent of ribosome | 16.5 | 1 | 9 |
| O25095 | ATP binding, tetraacyldisaccharide 4'-kinase activity | 35.5 | 1 | 5 |
| O25434 | 4 iron, 4 sulfur cluster binding, aspartic acid methylthiotransferase activity, metal ion binding, protein methylthiotransferase activity | 49.6 | 1 | 8 |
| O26001 | ATP binding, ATP hydrolysis activity | 29.2 | 1 | 14 |
| O25992 | Hypothetical protein | 82.3 | 1 | 3 |
| P56081 | 3-dehydroquinate synthase activity, metal ion binding, nucleotide binding | 39.1 | 1 | 3 |

### 3.7. Alterations in immune response due to immunization

The highest reciprocal titer of serum immunoglobulins was detected against outer membrane proteins (OMPs) and lipopolysaccharide (LPS) isolated from A61C (1). A sharp rise in the serum titer was observed against OMPs. Antibody titers were also detected in nonimmunized mice, but the titer was below the detection limits (**Figure 7**).

**Fig. 7:**
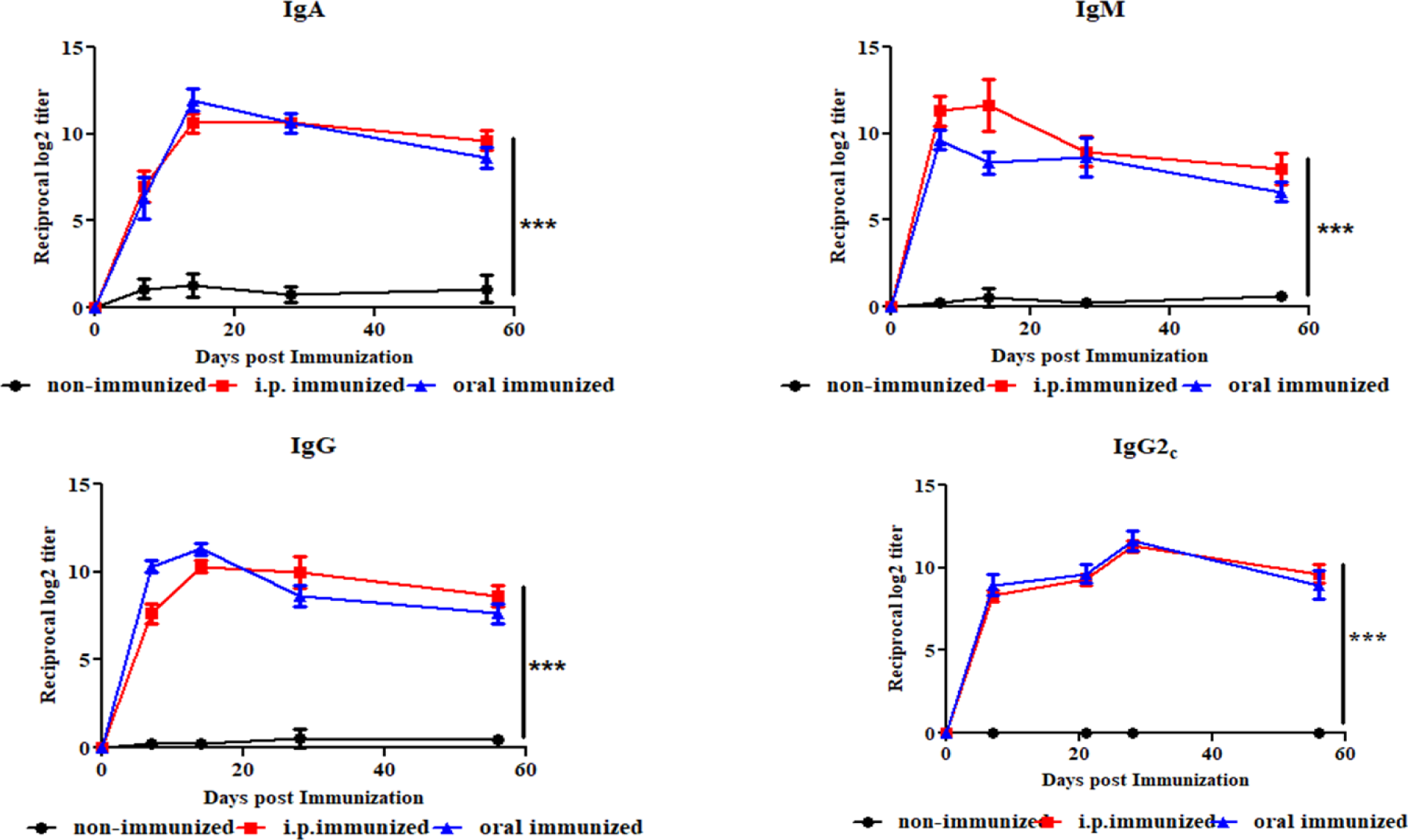
Elevation of immunoglobulins post immunization against OMPs. Data represented here are the mean values +/-Standard Deviations (SD) of three independent experiments. The differences in postimmunization day wise response of each of the studied cytokines were highly significant. Statistical significance was found between nonimmunized, i.p. and oral immunization mice serum (***p<0.001)

Spleen cells primed with OMV-based immunogens showed significant increases in IFN-γ, IL-1β, and IL-17 expression in the orally immunized groups compared with the i.p. immunized groups. No significant response was observed for IL-10, IL-4 and IL-6. TNF-α was elevated in the i.p. immunized mice than orally immunized (**Figure 8**).

**Fig. 8:**
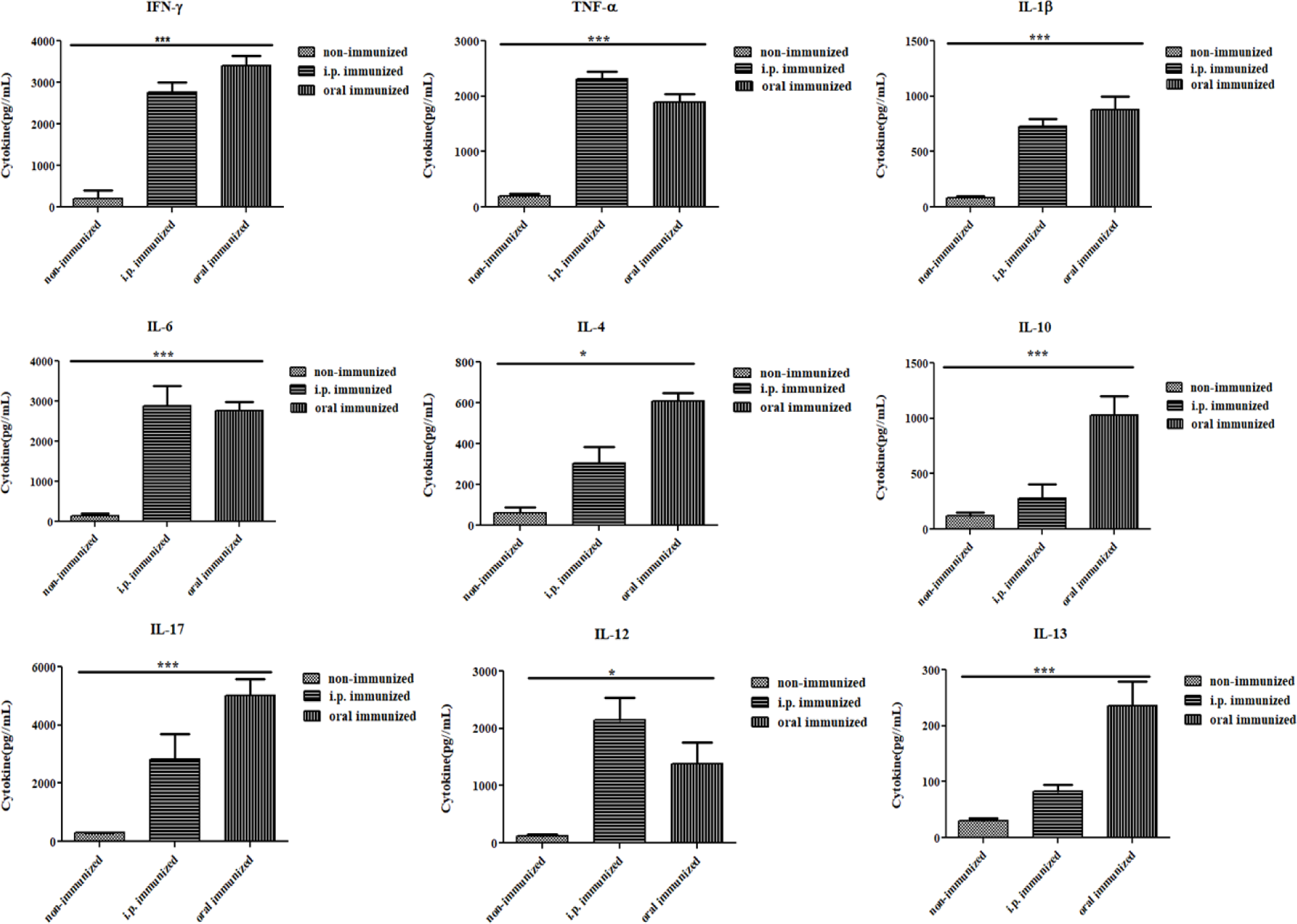
Elevation of inflammatory cytokines post immunization. Data represented here are the mean values +/-Standard Deviations (SD) of three independent experiments. The differences in postimmunization day wise response of each of the studied cytokines were highly significant. Statistical significance was found between unimmunized, i.p. and oral immunization mice serum (***p<0.001)

### 3.8. Serum Bactericidal Assay and Scanning Electron Microscopy (SEM)

The effect of immunized sera was shown by the absence of colonies compared to the control. Moreover, SEM images revealed that the bacterial surface of immunized serum-treated bacteria had been damaged, leading to death of the bacteria (**Figure 9**).

**Fig. 9:**
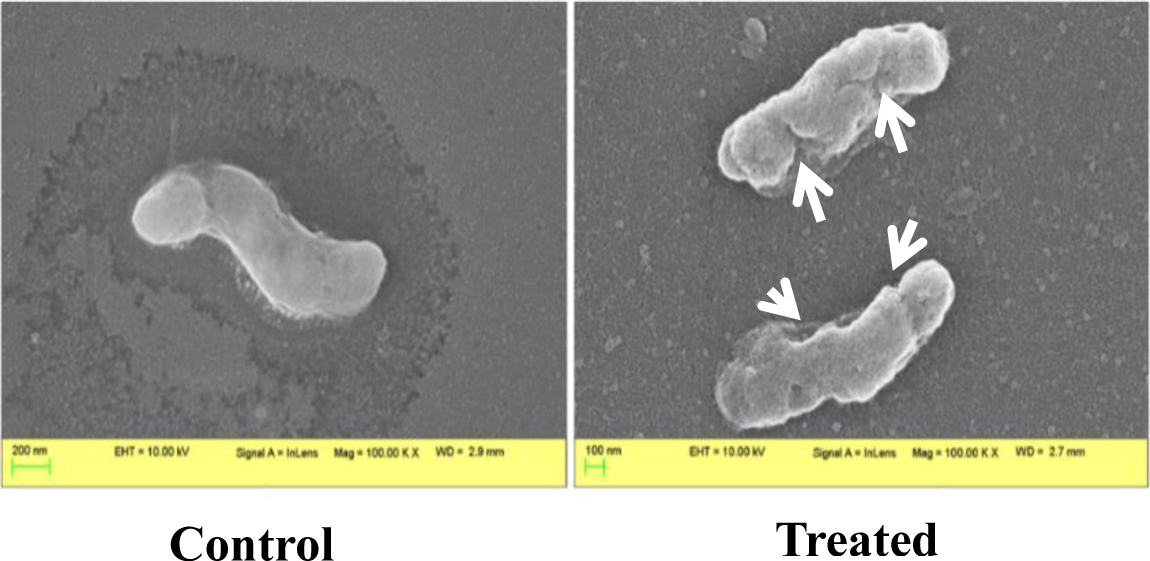
Serum Bactericidal Assay visualized under SEM.

### 3.9. Fluorescence-Activated Cell sorting analysis

Spleen cells tagged with anti-CD4+, anti-CD8+ and anti-CD19+ antibodies conjugated with FITC revealed a 30% increase in both the CD4+ and CD19+ populations and a 10% increase in the CD8+ population due to oral immunization, depicting the induction of possible T-cell or B-cell differentiation upon re-stimulation with OMVs (**Figure 10**).

**Fig 10.**
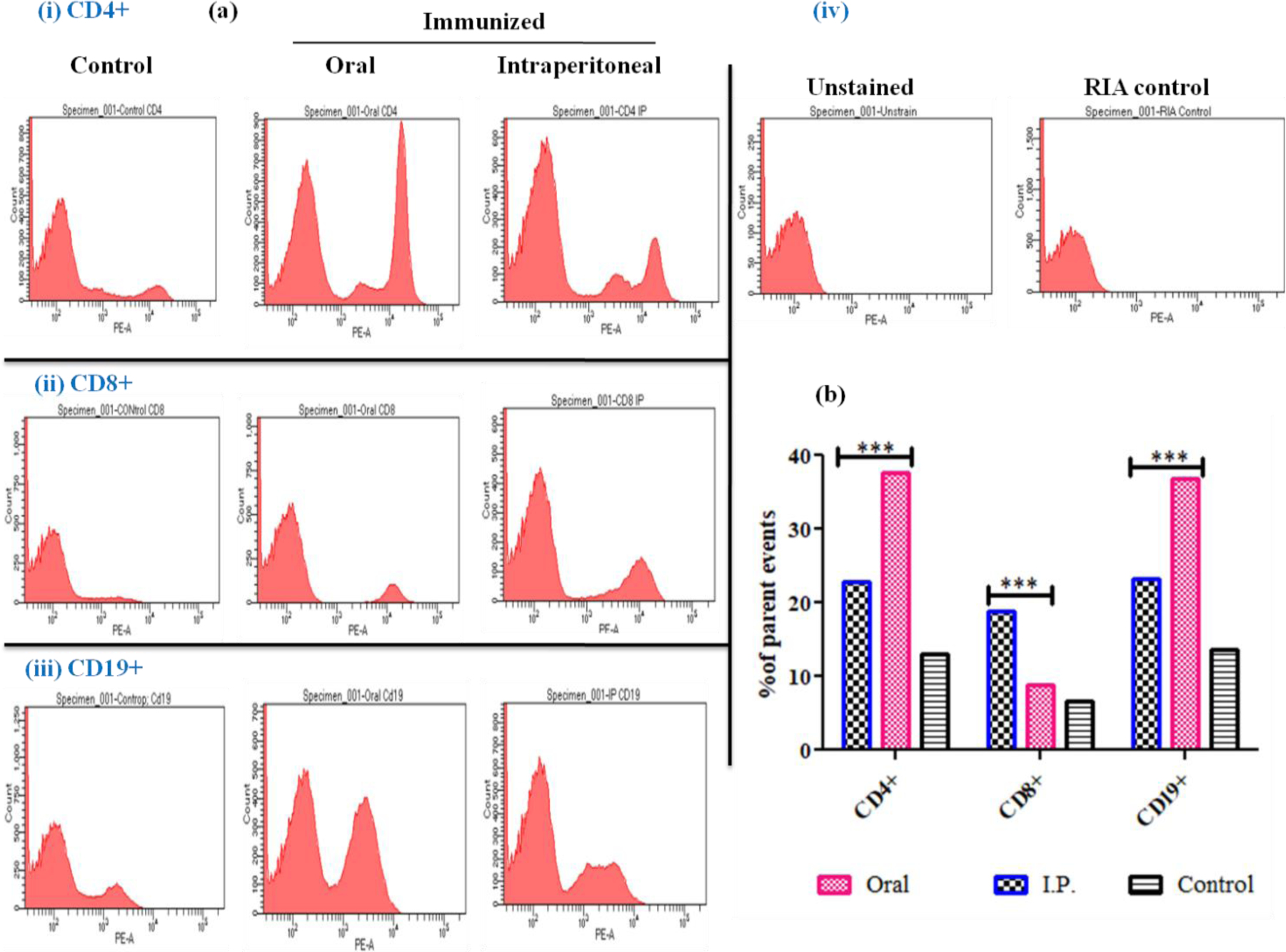
(a): Representative histograms of FACS analysis of OMVs immunized and nonimmunized CD 19+ (i), CD 4+ **(ii),** CD 8+ spleen cells **(iii)** stained using CD 19- PE, CD 4-PE, CD 8a-PE and counted in FACS Aria III flow-cytometer (BD Bioscience, USA) **(iv)** Represents unstained and RIA control. **(b) Bar diagram comparing the percentage of CD 19+, CD 4+, and CD 8a+ spleen cells from immunized and nonimmunized mice**. Two-way analysis of variance (ANOVA) test was used for statistical analysis. Bars represent mean ± S.E. of three individual experiments. Significant difference was found between OMVs immunized and nonimmunized spleen cell population (*** p value< 0.001).

### 3.10. Protective efficacy study post immunization

Nonimmunized mice showed extensive damage in terms of distortion in normal structure and inflammatory cell infiltration, whereas immunized mice showed significantly reduced inflammatory cell infiltration and preservation of structural integrity of the gastric epithelium.

Depending upon the severity of gastritis, a score was assigned, representing grade 0 being no change to grade 3 being severe gastritis. A reduction in gastric damage was observed in the immunized groups but not in the nonimmunized groups (**Figure 11**).

**Fig. 11:**
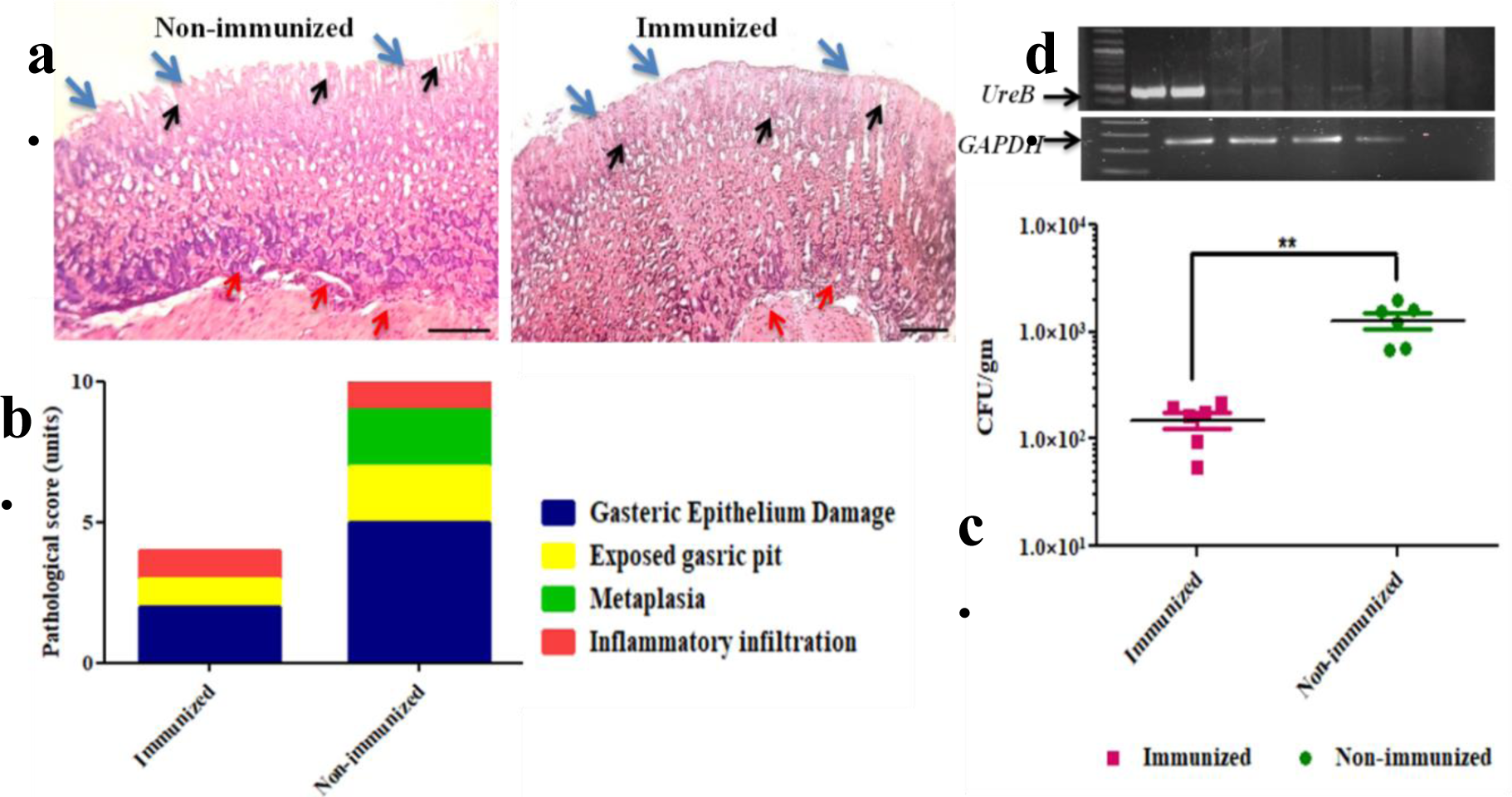
OMVs immunized mice serum reduces gastric tissue damage and inflammation in immunized mice after infection with SS1 (2x 10 CFU/mL). (a) Histological images represent both immunized and nonimmunized mice stomachs. OMVs immunized mice showed mild epithelial layer damage, less altered gastric mucosa and inflammatory infiltration, whereas nonimmunized mice displayed marked epithelial layer damage, inflammatory infiltration, exposed gastric pit and early signs of gastric metaplasia. (Blue arrow: gastric epithelium; Black arrow: exposed gastric pit; Red arrow: Inflammatory infiltration **(b)** Pathological scores of immunized or nonimmunized mice post challenge. **(c)** Tissue colonization, **(d)** Tissue DNA extraction and PCR amplification of ureB gene isolated from both immunized and nonimmunized mice. GAPDH represents as control for the mice.

## 4. Discussion

Over the years, a number of animal models have been evaluated for pathophysiology or treatment against *H. pylori,* including gnotobiotic pigs, dogs, cats, Mongolian gerbils, guinea pigs, rhesus monkeys, and mice [37–48]. In most cases, C57BL/6 or black mice were explored extensively because of their substantial contribution in *H. pylori-*related studies. The proper route for administering the pathogen and/or immunization also has a key impact on the development of an infection and assessment of immune response, which is another important consideration when selecting an animal model for *in vivo* studies. Therefore, cytokine alterations along with histological changes described in the present study represent a successful infection achieved through our newly developed surgical model. Conventional approaches for studying *H. pylori* infection in animals usually involve multiple oral inoculations using an oral gavage [49–51]. However, relying solely on the oral route to induce an infection and expecting the bacterium to outcompete the existing microflora and successfully colonize the stomach may not always yield a consistent result in any given experimental setting. This can cost significant time and resources being invested while still fostering uncertainty about an actual infective status in experimental animals. Therefore, it is important to consider the limitations and variability of the *in vivo* systems and look for alternative approaches that could provide a more reliable method of *H. pylori*- mediated pathogenesis in animal models.

Clinical detection of *H. pylori* infection generally involves histology and PCR apart from the Rapid Urease Test (RUT) [52–54]. Serological tests are often avoided, as previously invoked antibodies fail to recognize the actual infective status of recent manifestations [55–56]. As a consequence, this increases the chances of false positive results. In addition, histology allows visualization of pathogen-induced changes in gastric tissues, such as the intensity of inflammatory cell infiltration or aberrations in gastric topology, while PCR detects the presence of genomic DNA of *H. pylori* in gastric tissue samples [57]. However, it should be noted that neither histological observations nor negative PCR results rule out the absence of an infection [58]. Thus, a number of different techniques must be employed simultaneously to achieve a more accurate diagnosis of *H. pylori* infection [59]. Our study comprised a combination of histological observation, PCR detection and quantification of serum cytokine levels to confirm active *H. pylori* infection.

Surgical intervention initially spiked pro-inflammatory cytokines such as IFN-γ and IL-1β along with IL-17 significantly more on day 7 than on day 14. However, as the infection progressed, these cytokines were lowered and finally balanced, except for IL-6, which was found to be elevated more on day 14 than on day 7. A pronounced IL-6 level at later stages might indicate ongoing inflammation in the gastric lining with potentially developing chronic gastritis. Such responses were further supported by the induction of other cytokines. IFN-γ is an early effector molecule responsible for generating a Th1-mediated response by initiating different signaling cascades. However, upregulation of IFN-γ transiently downregulates IL-1β production. In addition, IL-17, a cytokine regulating the Th-17-based response, plays important roles in both pathogenesis and host immunity. Studies with chronic diseases have revealed that well-balanced IL-1β and IL-17 levels are constitutively produced to sustain inflammation due to infection in the long term. In the case of *H. pylori* infection, both IL-1β and IL-17 play crucial roles in pathogenesis; in particular, IL-17 influences the disease outcome upon infection. Our model showed an initial elevation in these cytokines, which decreased over time, indicating progression toward a chronic infection. However, as 7 days were not sufficient to develop a chronic infection, our model showed promising results in a time-dependent manner. Consistent with the cytokine analysis, histopathological observations also validate such changes to some extent. Intense inflammatory cell infiltration was observed on day 7 than on day 14, and the gastric lining was found to be more damaged with exposed gastric pits, indicating destruction caused by bacteria. Nevertheless, we did not find any striking structural abnormalities in gastric tissue 7 days post infection. PCR results from the same samples confirmed the presence of bacterial genomic DNA in experimental animals.

Next, we evaluated the surgical model for vaccine efficacy studies. Two different immunization routes were assessed to observe any alterations in the immune response due to changes in the route of administration. Immunization was performed both orally and intraperitoneally (i.p.) on days 0, 14 and 28. Initially, an elevation of serum IgG, IgM and IgA levels was observed against OMPs but not LPS of *H. pylori*. This can be due to the structural similarity between *H. pylori* LPS and blood antigens of the host [60–61]. Furthermore, we evaluated IgG2c (IgG subtype) and found it to be increased in immunized rather than nonimmunized groups [62]. Our study found oral immunization to be better responsive than i.p. route, which can be due to the presence of different surface proteins on OMVs that are more readily absorbed and reactive to gastric epithelial cells than peritoneal immune cells. A splenic cell re-stimulation (ex vivo) assay revealed enhanced Th2- based cytokine responses, such as IL-4, IL-13, IL-10 and IL-12, coinciding with previous studies with *H.* pylori-derived OMVs used as immunogens [63]. Interestingly, our study did not find any biased immune response against OMVs, indicating that the immune response to OMVs is not general but rather unique to each strain. CD4+, CD8+ and CD19+ cell populations were increased due to OMV immunization independent of the route of administration. OMV immunization ultimately leads to a reduction in bacterial colonization in immunized animals but not in nonimmunized animals. The serum bactericidal assay results showed that our immunized animal sera killed *H. pylori in the* presence of complement compared to the nonimmunized animal sera (31, 64, 65).

## 5. Conclusion

In conclusion, the intragastric surgical model of *H. pylori* infection can be used to study the pathophysiology, immune response, and potential therapies for *H. pylori* infection. Our study indicates that a minimum of 7 days is enough to develop an infection in this model. All experimental results showed that tissue samples collected at 7 days post infection can provide better results for diagnosing *H. pylori* infection than samples obtained at 14 days post infection, as histological changes and inflammatory cell infiltration are typically more pronounced at earlier time points post infection. Moreover, the cytokine response and antibody generation further support this model for vaccine efficacy studies. The immunization of mice with *H. pylori* OMVs has been shown to reduce the bacterial load with elevated antibody titers and protect gastric tissue from destruction. Therefore, the intragastric surgical model can become a valuable tool for understanding the pathophysiology of *H. pylori* infection, formulation and evaluation of potent vaccine candidates and development of potential therapeutics.

## 6. Author contributions

**Sanjib Das:** Conceptualization, experimental design and performance, data analysis and interpretation, manuscript preparation, **Prolay Halder:** Experiment performed, data analysis and interpretation, review and edit manuscript, **Soumalya Banerjee:** Experiment performed, data analysis and interpretation, review and edit manuscript, **Asish Kumar Mukhopadhyay:** Review and edit manuscript**, Shanta Dutta:** Review and edit manuscript, **Hemanta Koley**: Conceptualization, experimental design and supervision, data analysis and interpretation, manuscript preparation, funding acquisition.

## Acknowledgments

Our heartfelt thanks go to Mr. Suhashit Ranjan Ghosh, Mr. Subrata Singha, Mr. Pritam Nandy, and Mrs. Arpita Sarbajana for their technical assistance from time to time. We extend our gratitude to the University Grants Commission (UGC), New Delhi, India, for providing the fellowships to Sanjib Das under the CSIR-UGC-NFSC scheme (Student ID: SANJIB DAS under UGC-NFSC scheme 3363/(CSIR-UGC NET JUNE 2018]), the Indian Council of Medical Research for providing the fellowship to Prolay Halder [Fellowship ID: ICMR-3/1/3/JRF-2018/HRD- 066(66125)] and Soumalya Banerjee under the CSIR-UGC-NET scheme [Student ID:191620007740].

---

**Supplementary 1:** Antibiotic profile.

| Strains | Antibiotics |  |  |
| --- | --- | --- | --- |
|  | Clarithromycin | Amoxicillin | Metronidazole |

|  |  |  | S | R | S | R | S | R |
| --- | --- | --- | --- | --- | --- | --- | --- | --- |
| 1 | Ref. strains | 26695 | + | - | + | - | - | + |
|  |  | J99 | + | - | + | - | - | + |
|  |  | SS1 | + | - | + | - | - | + |
| 2 | BHU 8A |  | - | + | + | - | - | + |
| 3 | KO 8A |  | + | - | + | - | - | + |
| 4 | AS 2 |  | + | - | + | - | - | + |
| 5 | OT-10 (A) |  | + | - | + | - | - | + |
| 6 | B34 |  | + | - | + | - | - | + |
| 7 | D383 |  | + | - | + | - | - | + |
| 8 | B6 |  | + | - | + | - | - | + |
| 9 | M28 |  | + | - | + | - | - | + |
| 10 | L7 |  | + | - | + | - | - | + |
| 11 | A61C(1) |  | + | - | + | - | - | + |
| 12 | AM1 |  | + | - | + | - | - | + |
| S - sensitive |  |  |  |  | R – resistant |  |  |  |

**Supplementary 2:**
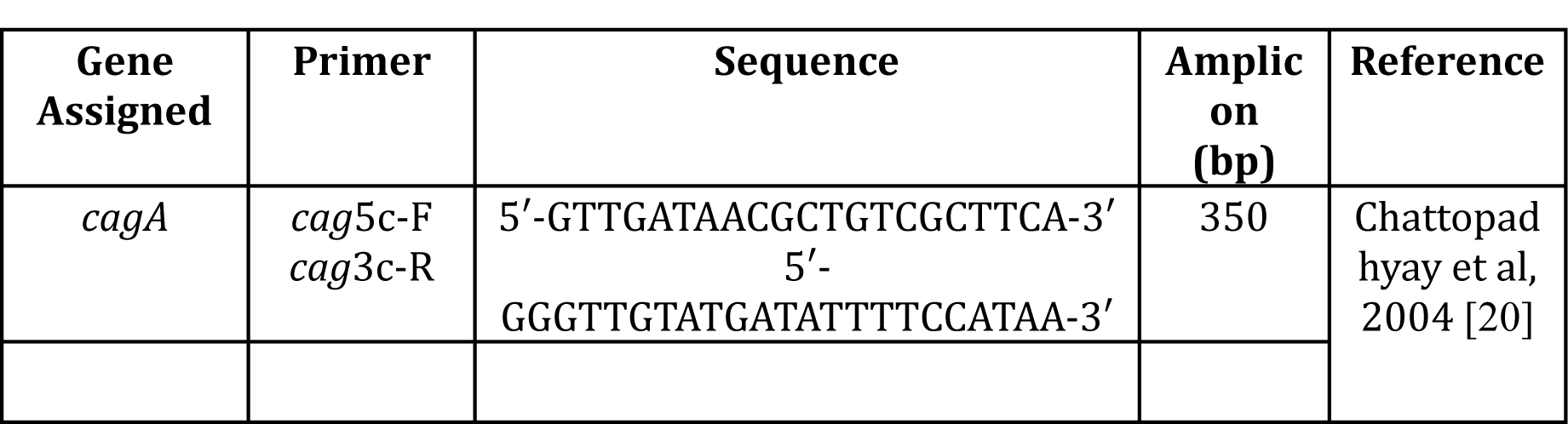
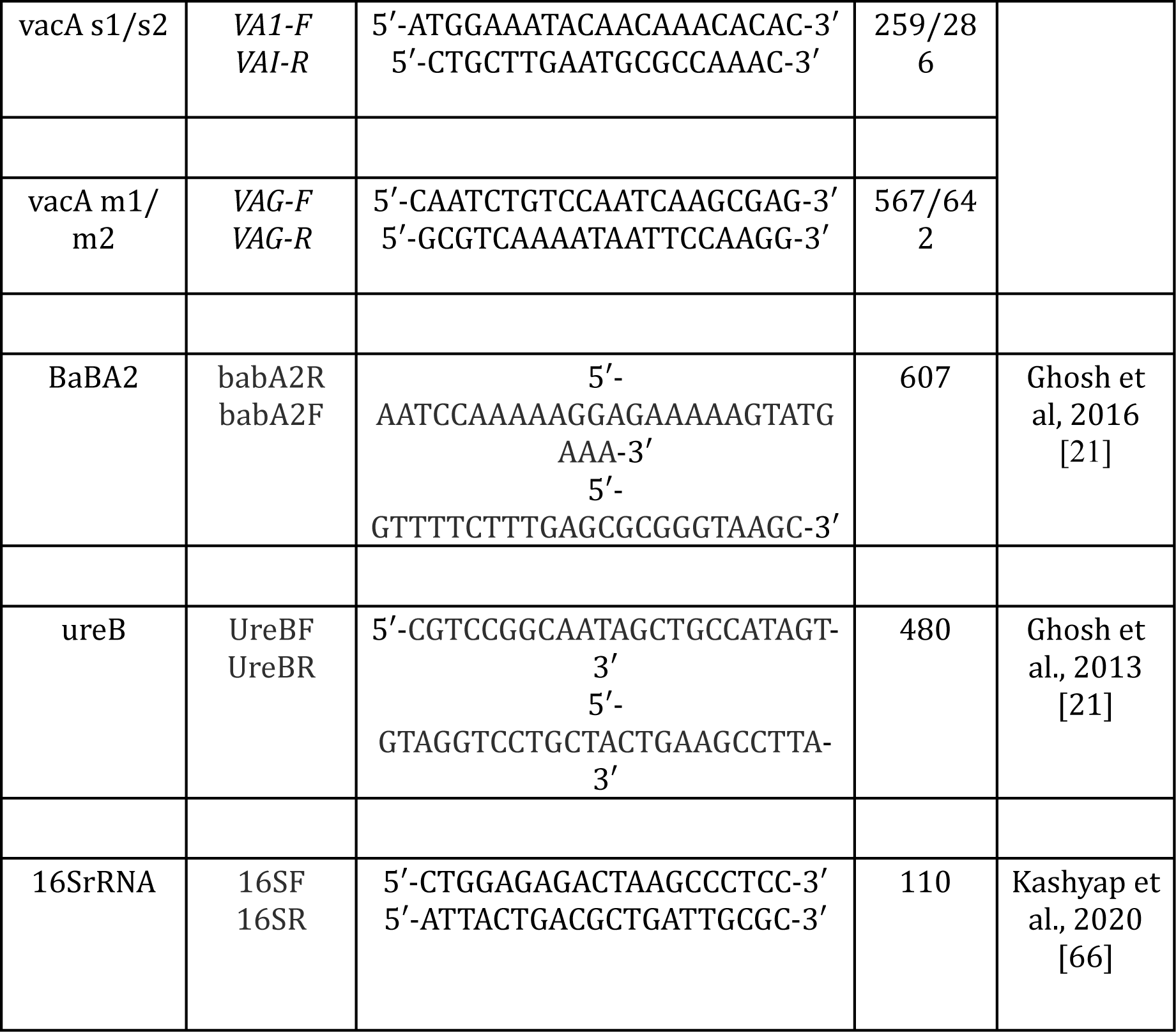
Primers used in this study.

